# Identification and characterization of long non-coding RNAs as targets of mammary tumor cell proliferation and migration

**DOI:** 10.1101/036418

**Authors:** Sarah D. Diermeier, Kung-Chi Chang, Susan M. Freier, Junyan Song, Alexander Krasnitz, Frank Rigo, C. Frank Bennett, David Spector

## Abstract

Recent genome-wide studies revealed that as much as 80% of the human genome can be transcribed whereas only 2% of this RNA is translated into proteins. Non-coding transcripts can be subdivided into several groups, with long non-coding RNAs (lncRNAs) representing the largest and most diverse class. With breast cancer being the most frequent malignancy in women worldwide, we set out to investigate the potential of lncRNAs as novel therapeutic targets. By performing RNA-Seq on tumor sections and mammary organoids from MMTV-PyMT and MMTV-Neu-NDL mice, modeling the luminal B and HER2/neu-amplified subtypes of human breast cancer respectively, we generated a comprehensive catalog of differentially expressed lncRNAs. We identified several hundred potentially oncogenic lncRNAs that were over-expressed in a subtype-specific manner as well as numerous lncRNAs up-regulated in both models. Among these lncRNA we defined a subset of 30 previously uncharacterized lncRNAs as Mammary Tumor Associated RNAs (MaTARs) and we identified human orthologs. We functionally validated the role of these MaTARs by antisense oligonucleotide (ASO) mediated knockdown in primary mammary tumor cells and 3D *ex vivo* organoids. Upon independent knockdown of 15 MaTARs, we observed significantly reduced cell proliferation, invasion and/or collective cell migration in a cancer-specific context. Thus, MaTARs are likely key drivers of mammary tumor progression and/or metastasis and represent promising new therapeutic targets.

## Reviewer link to deposited data

http://www.ncbi.nlm.nih.gov/geo/query/acc.cgi?token=wnqvokgoxtaxdod&acc=GSE72823

## Introduction

Breast cancer is the most frequent malignancy in women worldwide and the second leading cause of cancer mortality in women (Siegel et al. 2015). The challenge of finding more efficient treatments is complicated by the diversity of the disease, resulting in the classification of numerous breast cancer subtypes. The most common type, invasive ductal carcinoma, can be stratified into two fundamental classes: the estrogen receptor (ER) and/or progesterone receptor (PR) positive subtypes (luminal A and luminal B) and ER-negative (HER2/neu-amplified and basal-like) disease. These "intrinsic" breast cancer subtypes differ in gene expression patterns, histo-pathology, patient prognosis and treatment strategy (Perou et al. 2000; Sorlie et al. 2003; Cancer Genome Atlas Network 2012; Anderson et al. 2014). More recently, 10 subgroups of breast cancer were proposed based on genome-wide copy number variations and expression studies (Curtis et al. 2012; Ali et al. 2014).

While currently available endocrine and targeted therapies have lead to improved overall survival rates, many breast tumors are intrinsically resistant or acquire resistance after initial responsiveness (for review, see Ali and Coombes 2002; Higgins and Baselga 2011). To improve the existing treatment regimes, it is critical to identify and investigate new molecular targets that have the potential to inhibit breast cancer progression and metastasis by impacting alternative pathways and/or restoring drug sensitivity. Thus far, all efforts of drug development have focused on proteins, such as ER, HER2/neu, aromatase, the mammalian target of rapamycin (mTOR), phosphatidylinositol 3-kinase (PI3K), protein kinase b (Akt/PKB), cyclin-dependent kinases (CDKs), growth factors or histone deacetylases (HDACs) (for review, see Yamamoto-Ibusuki et al. 2015).

The ENCODE consortium revealed that only 2% of all genes have the ability to be translated into proteins, while as much as 80% of the human genome can be transcribed in a cell type specific manner (Okazaki et al. 2002; Bertone et al. 2004; Katayama et al. 2005; Carninci et al. 2005; Kapranov et al. 2007; Djebali et al. 2012; ENCODE Project Consortium 2012). Non-coding transcripts are subdivided into several classes, with long non-coding RNAs (lncRNAs) representing the largest and most diverse class. They are defined by length, ranging from 200 nucleotides to 100 kilobases. Like mRNAs, they are transcribed by RNA polymerase II and can be capped, spliced and poly-adenylated. However, lncRNAs lack a significant open reading frame, can be transcribed from the sense or antisense orientation and commonly originate from introns or intergenic regions (Derrien et al. 2012; St Laurent et al. 2012; Iyer et al. 2015). Members of this class of non-coding transcripts have been implicated as regulatory molecules in a variety of cellular functions (for review, see Rinn and Chang 2012; Kornienko et al. 2013; Bergmann and Spector 2014).

LncRNAs represent a relatively unexplored class of regulatory molecules to pursue as potential therapeutic targets (for review, see Prensner and Chinnaiyan 2011; Wapinski and Chang 2011; Shore et al. 2012; Wahlestedt 2013; Cheetham et al. 2013). One of the few examples of lncRNAs that have been studied in the context of breast cancer is the HOX antisense intergenic RNA (HOTAIR) (Rinn et al. 2007). It promotes mammary tumor invasion and metastasis and acts as an independent predictor of patient survival rates (Rinn et al. 2007; Gupta et al. 2010). Elevated expression levels of another lncRNA BCAR4 were described to correlate with a higher rate of metastasis and shorter patient survival (Xing et al. 2014). Recently, lncRNA152 and lncRNA67 were shown to play a role in the growth of breast cancer cell lines (Sun et al. 2015). Eleanors, a new class of lncRNAs, were found to be associated with breast cancer resistance and adaptation (Tomita et al. 2015). Differential expression of a few additional lncRNAs has been studied in the context of breast cancer, but as for most lncRNAs the functional mechanisms remain elusive (for review, see Hansji et al. 2014; Vikram et al. 2014). As only a handful of the 16,000 annotated lncRNAs (GENCODE v22) have been functionally characterized and studied in the context of breast cancer, it is essential to perform an unbiased RNA-Seq screen to identify the complement of lncRNAs that exhibit altered expression in mammary tumors and as such represent potential therapeutic targets.

Here, we generated a comprehensive catalog of lncRNAs that are up-regulated in primary mammary tumors compared to the normal mammary gland epithelium. We performed RNA-Seq analysis on three physiologically relevant transgenic mouse models of luminal B (MMTV-PyMT) and HER2/neu-amplified (MMTV-Neu-NDL and MMTV-Cre;Flox-Neo-Neu-NT) subtypes of breast cancer (Guy et al. 1992; Siegel et al. 1999; Andrechek et al. 2000). Among the 290 lncRNAs that are upregulated at least 2-fold in the tumors compared to normal mammary glands we prioritized 30 previously uncharacterized lncRNAs as <u>Ma</u>mmary <u>T</u>umor <u>A</u>ssociated <u>R</u>NAs (MaTARs). In addition, we identified a gene coexpression network that is highly specific for the mammary epithelium. In order to functionally validate these MaTARs as key drivers in tumor progression, we performed antisense knockdown studies in primary cancer cells and 3D *ex vivo* mammary organoids. Our results indicate that several of the investigated MaTARs are involved in mammary tumor cell proliferation and/or collective cell migration and as such represent exciting candidate therapeutic targets.

## Results

### Comparing the transcriptome of mammary tumors and corresponding organoids

Recently, the culturing of mammary organoids in 3D artificial extracellular matrix (ECM) hydrogels has gained popularity over 2D cell culture approaches, especially for studying mammalian development and disease (for review, see Shamir and Ewald 2014). Previous studies demonstrated that mammary branching morphogenesis can be recapitulated in an organoid system by retaining its epithelial spatial organization (Barcellos-Hoff et al. 1989; Fata et al. 2007; Ewald et al. 2008; Nguyen-Ngoc et al. 2012). Furthermore, organoids that closely resemble tumors have the potential to serve as a rapid *in vitro* screening system in drug development and "personalized medicine". However, thus far the transcriptome of organoids has not been examined to determine how closely they resemble the gene expression signature of the original tissue they were derived from.

In this regard, we removed mammary tumors of comparable volume, histological profile, and expression of several breast cancer markers (ER, PR and HER2) from three MMTV-PyMT mice (Figure S1A). For each tumor, one half was frozen, cryosectioned and RNA was isolated directly from the tissue sections (Figure 1A). The other half of the tumor was used to generate tissue organoids and RNA was extracted either immediately following organoid preparation (day 0) or after 6 days of culturing in 3D matrigel domes (day 6).

**Figure 1.**
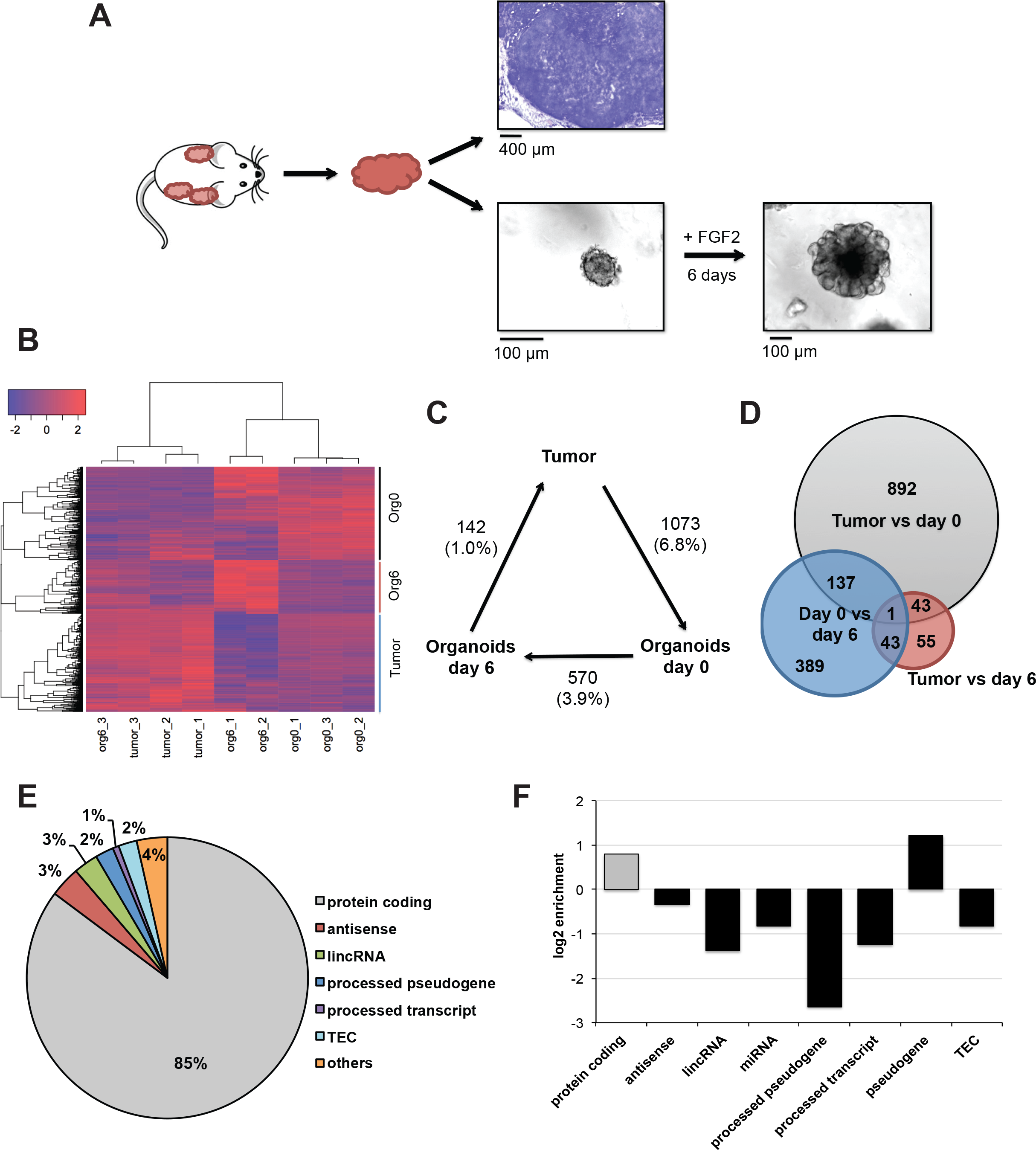
Comparison of tumor and organoid transcriptomes. A) Tumors isolated from three MMTV-PyMT mice were cut in half. One half was cryosectioned (30 μm sections) and RNA was isolated directly from the tissue. Toluidine Blue staining of a representative section (10 μm) is shown (upper panel). The other half of the tumor was used to generate organoids and RNA was isolated either immediately (day 0, left lower panel) or after 6 days of culturing in the presence of FGF2 (day 6, right lower panel). B) Variance-stabilizing transformed (vst) read counts of genes that were significantly (FDR < 0.1) differentially expressed in the tumor, organoids day 0 and/or organoids day 6 datasets (total number of genes across all three datasets: 1560). Hierarchical clustering identified three clusters (Org0, Org6 and Tumor) that are defining the respective datasets. Genes within the "Org0" cluster are up-regulated during organoid preparation but reverted back to the levels in tumors after 6 days in culture. Counts are scaled per row, hence "0" indicates no change between tumor, organoids day 0 and organoids day 6. Red denotes "higher expression levels" and blue indicates "lower expression levels" compared to the other two groups. C) Illustration of the number of genes significantly altered comparing tumor, day 0 and day 6 datasets. Percentage of the transcriptome is denoted in brackets, i.e. 142 genes represent 1.0% of the transcriptome in day 6 organoids compared to tumor sections. D) Venn diagram indicating the overlap of differentially expressed genes across all datasets. E) Transcript biotypes of the 142 differentially expressed genes comparing day 6 organoids to tumors. The majority (85%) of altered transcripts are protein-coding. TEC = To be Experimentally Confirmed (GENCODE classification). F) Log2 enrichment of transcript biotypes in E), normalized to the genome-wide frequency of these biotypes. Protein-coding genes and pseudogenes are overrepresented within the dataset, whereas most non-coding biotypes are underrepresented.

Comparative RNA-Seq analyses revealed that the global gene expression signature of mammary tumor organoids correlates well with the tumor transcriptome (Pearson correlation coefficient = 0.82 for day 0 and 0.84 for day 6, see Figure S1B-C). We observed significant expression changes for 1073 genes (or 6.8% of the transcriptome, Figure 1B-D) when comparing the tumor sections to organoids at day 0. The "Org0" cluster in Figure 1B indicates a subset of genes that are up-regulated in organoids at day 0 compared to the tumor transcriptome. Interestingly, the expression levels of many of these genes were subsequently down-regulated in the course of the culturing period (day 6), representing the original expression level in the tumor ("Org0" cluster in Figure 1B, general direction of altered gene expression in Figure S1D, S2A). We conclude that the transcriptome undergoes temporary changes during the process of organoid preparation due to a stress response. These changes are subsequently reverted as the organoids "recover" during the culturing period.

Importantly, only 1% of the transcriptome (142 out of the 13,854 expressed genes) showed statistically significant alterations (Figure 1B-D, S1D) between the tumor and the day 6 organoids. The affected genes were almost exclusively (85%) protein coding genes (Figure 1E). In comparison, only 48% of all genes in the most recent GENCODE annotation vM5 are classified as "protein coding". Hence, the subset of genes that was significantly altered in our dataset is enriched for protein coding genes and depleted of non-coding RNA species except pseudogenes when compared to the global abundance of these transcript biotypes (Figure 1F). While we observe significant gene expression changes for only 142 genes, we note that several pathways involved in signaling and metabolism appeared altered when taking into account genome-wide fluctuations (Figure S2B). Thus, the first direct transcriptomic comparison of cultured organoids and the tumors they were derived from demonstrates that the gene expression signature of day 6 mammary organoids closely resembles the tumor transcriptome and represents a reliable model system to study the expression of non-coding RNAs.

### The transcriptome of luminal B and HER2-amplified mammary tumors

To identify the complement of lncRNAs expressed in different subtypes of mammary carcinomas, we generated organoids from two MMTV-PyMT and three MMTV-Neu-NDL mice. After 6 days in matrigel, the transcriptome of the organoids was analyzed using RNA-Seq. To classify cancer-specific changes in the gene expression signature in comparison to normal tissue, we also generated and sequenced organoids from nulliparous mammary glands of three age-matched wild type FVB females (Figure 2A).

**Figure 2.**
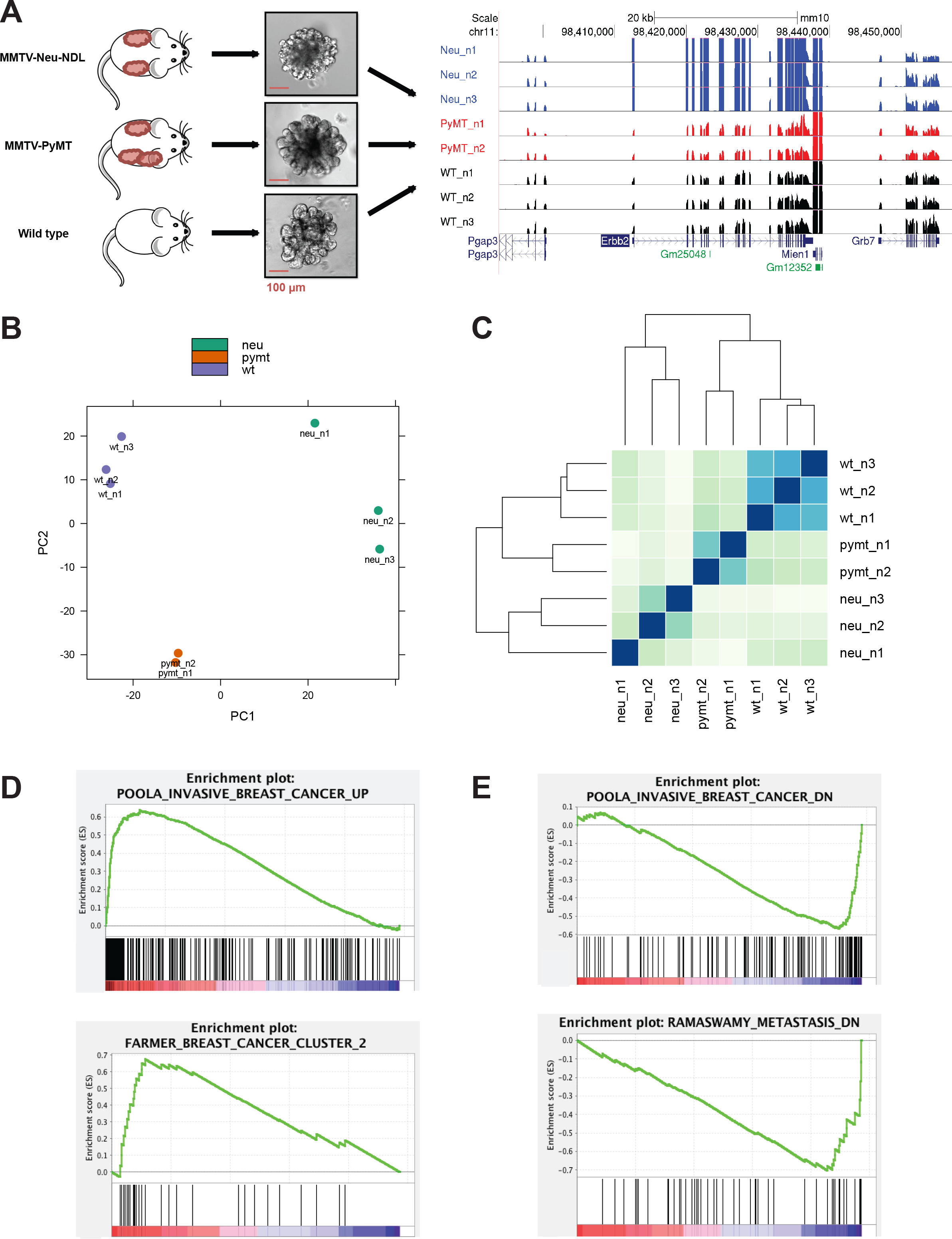
The transcriptomes of MMTV-PyMT and MMTV-Neu-NDL tumors. A) Schematic representation of the comparative RNA-Seq performed. Organoids were prepared from MMTV-PyMT, MMTV-Neu-NDL and wild type normal mammary glands and cultured for 6 days in matrigel (left). UCSC genome browser image (right) displays bedgraphs of RNA-Seq data exemplarily at the *Erbb2/neu* locus. Vertical viewing range: 0-50. B) Principal component analysis comparing the transcriptome of organoids derived from WT (purple), MMTV-PyMT tumors (orange) and MMTV-Neu-NDL tumors (green). Three clusters were detected according to the three different mouse models. C) Distance dendrogram of the transcriptome of organoids derived from WT (wt), MMTV-PyMT tumors (pymt) and MMTV-Neu-NDL tumors (neu). D) Examples of gene set enrichment analysis (GSEA) datasets found up-regulated in MMTV-Neu-NDL tumors (upper panel) and MMTV-PyMT tumors (lower panel). FDR < 0.01. E) Examples of gene set enrichment analysis (GSEA) datasets found down-regulated in MMTV-Neu-NDL tumors (upper panel) and both MMTV-Neu-NDL and MMTV-PyMT tumors (lower panel). FDR < 0.01.

Our RNA-Seq screen revealed significant changes in both the coding and the long non-coding transcriptomes, with a total of 4,633 (35% of all expressed genes) genes differentially expressed in MMTV-PyMT-and 4,322 (28% of all expressed genes) in MMTV-Neu-NDL-derived organoids (Figure S3A). As an internal control, we monitored the expression of *Erbb2/neu,* the driving oncogene in mouse models resembling the HER2/neu-amplified subtype of breast cancer. As expected, *Erbb2/neu* was overexpressed only in the MMTV-Neu-NDL organoids but not in the PyMT or wild type (WT) organoids (Figure 2A, S3B). Principal component analysis and Euclidean distance plots comparing WT mammary glands to the luminal B and HER2/neu-dependent mammary tumors revealed distinct clustering according to the tumor-driving transgene (Figure 2B, C).

Furthermore, we observed general and subtype-specific expression changes in cellular pathways as well as gene ontology (GO) terms (Figures S4, S5). Both subtypes exhibited an up-regulation of the tumor necrosis factor (TNF) signaling pathway, an indicator of antitumoral response of the immune system (Wajant et al. 2003). Subtype-specific up-regulation of "G-protein coupled receptor activity" was observed in the MMTV-Neu-NDL organoids (Figure S5A). The driving oncogene *Erbb2/neu* is a receptor tyrosine kinase (RTK) (Akiyama et al. 1986; Stern et al. 1986) and cross-talk between this receptor and G-protein couple receptors has been previously reported (Negro et al. 2006). In the MMTV-PyMT dataset, "cell cycle" was one of the most distinctively up-regulated pathways (Figure S4B) while "cell adhesion molecules" was down-regulated. In addition, we observed enrichment for the GO terms "cell migration", "production of cytokines", "inflammatory response" and "positive regulation of angiogenesis" when analyzing the up-regulated genes in MMTV-PyMT organoids (Figure S5B). These GO terms indicate the high proliferative rate and invasiveness commonly observed in breast cancer, as well as the inflammation and neo-angiogenesis at the site of the tumors (for review, see Hanahan and Weinberg 2011).

Importantly, gene-set enrichment analyses (GSEA) revealed that our mouse mammary organoid data resembled the transcriptome of human breast cancer. For instance, the down-regulated genes of both models correlated well with expression signatures of invasive breast cancer and metastatic tumors. A similar correlation was found for up-regulated genes in both datasets (Figure 2D - E). Taken together, these data demonstrate that mouse mammary carcinoma-derived organoids recapitulate the transcriptome of luminal B and HER2/neu-amplified breast cancer.

### Identification of up-regulated lncRNAs in mammary organoids

We set out to generate a catalog of all currently annotated lncRNAs that exhibit altered expression levels in mammary tumor organoids to provide a useful resource for both basic research and target development. We identified 484 lncRNAs in MMTV-PyMT and 402 lncRNAs in MMTV-Neu-NDL with an overlap of 122 between both models that are differentially expressed in organoids derived from mammary tumors compared to WT mammary glands (Figure 3A). These genes represent a diverse group of lncRNAs in terms of their biotypes and genomic location (Figure S6A-B). Here, we define "lncRNA" as any long transcript that is not classified as "protein coding", including, but not limited to, pseudogene and antisense transcripts. Approximately 30% of the identified RNA genes are classified as "lincRNA" (long intergenic non-coding RNAs), 23% as "processed pseudogenes", 16% as "antisense" transcripts, and 13% as "processed transcripts." (Figure S6A).

**Figure 3.**
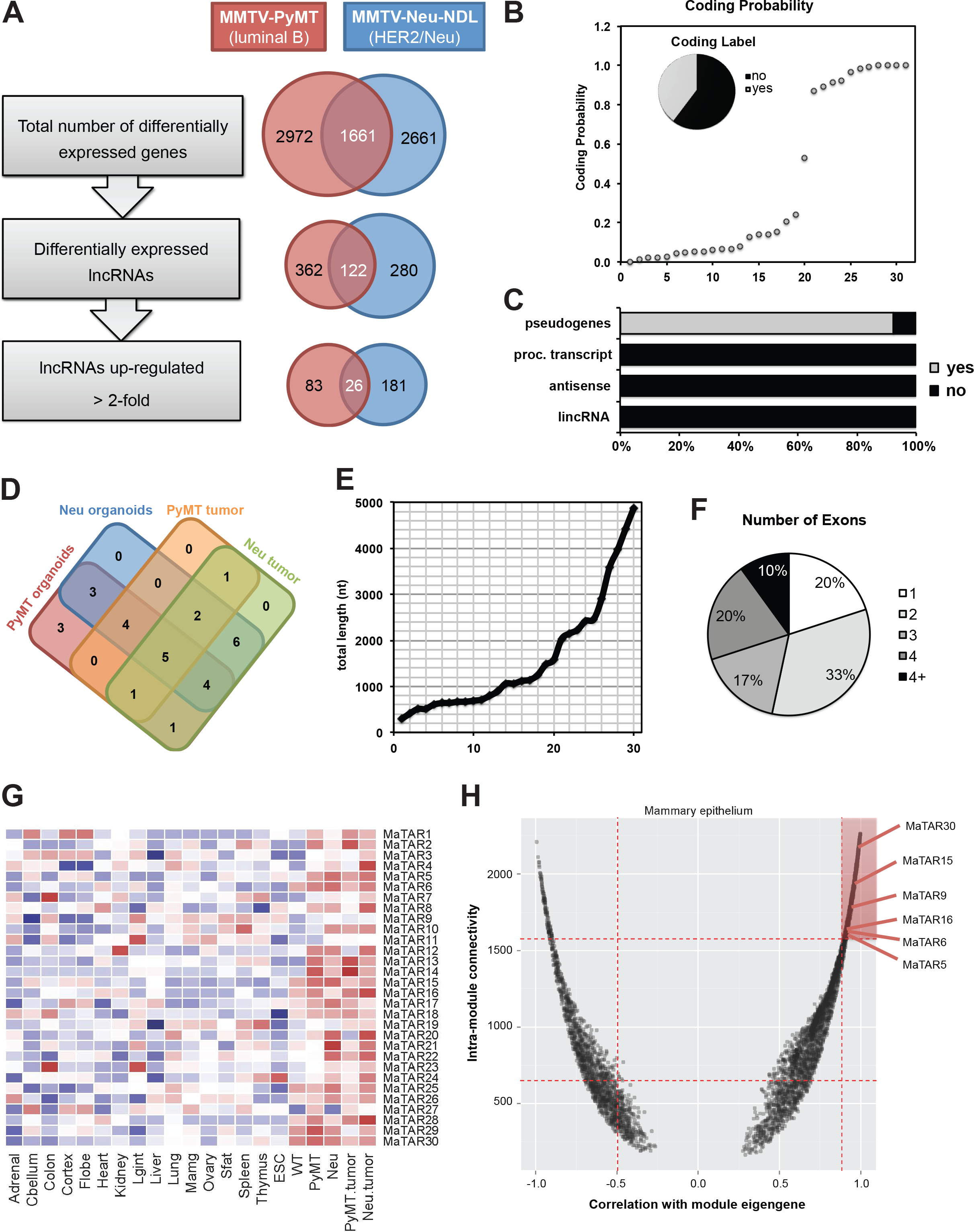
Identification and characterization of MaTARs. A) Computational analysis pipeline to identify potentially oncogenic lncRNAs. Numbers indicate genes significantly differentially expressed (adj. p-value < 0.1). Ensembl IDs and log2(fold-change) of all up-regulated (> 2-fold) genes is provided in Table S1 and S2. B) Coding probability of all 30 MaTARs according to CPAT. Coding probability is plotted on a scale from 0 to 1, with 0 indicating "no coding potential" and "1" representing "high coding potential". Insert: 63% of all MaTARs are non-coding (no) whereas 37% are potentially protein-coding (yes). C) Correlation of the CPAT coding label and lncRNA biotypes. The 37% of transcripts with protein-coding potential in B) are attributed to the pseudogene fraction of MaTARs. Not all pseudogenes are classified as protein coding. D) Venn diagram illustrating overexpression of the 30 MaTARs in four tumor models. PyMT tumor = tumor sections from MMTV-PyMT sections; Neu tumor = tumor sections from MMTV-Cre;Flox-Neo-Neu-NT. Most MaTARs are up-regulated in several datasets. Log2(fold-change) of all MaTARs in the four tumor models is provided in Table S3. E) Length distribution of all 30 MaTARs. The majority (20 MaTARs) is < 1500 nt long. F) Number of exons per MaTAR. More than 50% of MaTARs consist of one or two exons and only 3 transcripts have more than four exons. G) MaTAR expression (vst counts) across ENCODE tissues, wild type (WT) organoids and tumor organoids and sections. The highest expression level for 18 MaTARs was detected in tumors. All MaTARs are expressed in a tissue-specific manner. PyMT.tumor = tumor sections from MMTV-PyMT sections; Neu.tumor = tumor sections from MMTV-Cre;Flox-Neo-Neu-NT. Counts are scaled per row. H) Connectivity between genes in the "mammary epithelium" module is plotted against the module's eigengene correlation. Red dotted lines correspond to first and third quartiles. Six MaTARs are in the fourth quartile, denoted by high eigengene correlation and intra-module connectivity. Network modules of all MaTARs are described in Table S4.

Further, we aimed to focus on potentially oncogenic lncRNAs that might act as drivers of tumorigenesis and/or metastasis. Therefore, we filtered our dataset for lncRNAs that are up-regulated at least 2-fold compared to normal mammary organoids. We identified 109 up-regulated lncRNAs in MMTV-PyMT (Table S1) and 207 in MMTV-Neu-NDL organoids (Table S2), with an overlap of 26 lncRNAs (Figure 3A, highlighted in grey in Table S1 and S2). Interestingly, very few of the identified transcripts have been studied previously. One of these lncRNAs is Trp53cor1 (also known as lincRNA-p21), which is up-regulated at least 4-fold in both mammary tumor models (Table S1 and S2) and has been implicated in p53/p21-dependent gene regulation (Huarte et al. 2010; Dimitrova et al. 2014) as well as cancer (Yang et al. 2014; Chou et al. 2015). Furthermore, the two <u>Pl</u>uripotency-<u>a</u>ssociated <u>tr</u>anscripts Platr4 and Platr7 were up-regulated more than 4-fold in MMTV-Neu-NDL organoids (Table S2). Notably, Platr lncRNAs were identified in an RNA-seq screen of embryonic stem cells (ESCs) and shown to be correlated to the maintenance of the ESC expression profile (Bergmann et al. 2015). We performed coding potential analysis on all 290 lncRNA genes (443 associated transcripts in total) using the Coding-Potential Assessment Tool (CPAT) (Wang et al. 2013). Our results reveal no coding potential for 75% of the identified transcripts (Figure S6C-F) despite including pseudogenes in our analysis that typically score a high coding probability. As expected, we observe a good correlation between coding probability and ORF length but not RNA length (Figure S6E-F).

We further characterized the identified differentially expressed lncRNA genes regarding shared transcription factors binding to their promoter regions. Therefore, we performed motif analyses in the regions of −900 bp to +100 bp around the TSS of all up-regulated and all down-regulated non-coding RNA genes, identified in both PyMT - and Neu - derived organoids. For down-regulated promoters, we observe a 2-3-fold enrichment of a rather rare motif that matches Zbtb33/Kaiso as well as a 6-fold enrichment of motifs for NF-_K_B and CEBP (Figure S7A). 1. In contrast, up-regulated lncRNA genes seem to be primarily regulated by androgen receptor (AR) and p63 (Figure S7B).

To narrow down the long list of candidates to the ones with highest potential clinical relevance, we prioritized the transcripts further based on several stringent criteria: i) statistical significance (FDR adjusted p-value < 0.1), ii) at least 2-fold up-regulation in tumor organoids and/or tumor sections compared to normal mammary organoids, iii) sufficient read coverage per transcript to eliminate very lowly abundant RNAs (Figure S8A), iv) human conservation based on sequence and/or synteny, v) location in an intergenic genomic region and vi) lack of highly repetitive elements. The last two points ensure potential "druggability", i.e. that antisense oligonucleotides (ASOs) can be designed specifically for the lncRNA of interest. This is of high importance as we propose ASOs as a new treatment for breast cancer. Out of 290 up-regulated lncRNAs, 30 transcripts fulfill the above-mentioned criteria and were selected as an initial subset for further characterization. All 30 are overexpressed in at least 4 out of the 5 sequenced PyMT organoid datasets as well as significantly up-regulated in PyMT tumor sections (Figure 3D). We also included RNA-Seq data from MMTV-Cre;Flox-Neo-Neu-NT tumors, a second model for the HER2/neu-amplified subtype of human breast cancer, to further refine our candidate lncRNAs. We termed these transcripts <u>Ma</u>mmary <u>T</u>umor <u>A</u>ssociated <u>R</u>NA 1-30 (MaTAR1-30). A complete list of their official gene IDs and expression levels in all evaluated mammary tumor models can be found in Table S3. Of the 30 MaTARs, only MaTAR4 and MaTAR8 have been characterized in the past. MaTAR4 (Hoxa transcript antisense RNA, myeloid-specific 1; Hotairm1) has been implicated in myelopoiesis and myeloid maturation (Zhang et al. 2009; 2014). MaTAR8 (lncRNA-Smad7) has been described as a TGF-β - regulated antisense transcript of *Smad7* inhibiting apoptosis in mouse breast cancer cells (Arase et al. 2014). We tested the coding potential of all MaTARs using CPAT (Wang et al. 2013). While 60% of the MaTARs were classified as non-coding, 40% were predicted to be coding, attributed to the pseudogene fraction (Figure 3B-C). We further characterized the MaTARs according to their total length and number of exons (Figure 3E-F). MaTARs range in size from 291 - 4867 nt with the majority of MaTARs being shorter than 1500 nt. More than half of all MaTARs consist of one or two exons, only 10% show complex structures with more than 4 exons.

In order to compare the expression levels of MaTARs in mammary tumors to other tissue types, we analyzed the ENCODE expression datasets of adult mouse tissue as well as embryonic stem cells (Bergmann et al. 2015). We applied the same computational pipeline for mapping and read counting as was used for our tumor data and performed variance-stabilizing transformation (vst) of the resulting counts (Figure 3G). Our results confirm the general notion of tissue-specific expression of lncRNAs (Cabili et al. 2011; Derrien et al. 2012), with no MaTAR being expressed in high levels (vst counts > 10) in more than one tissue type. Interestingly, the majority of the MaTARs (18 out of 30) are expressed at very low levels in other tissues, indicating their therapeutic potential. MaTAR1, 3, 7, 9, 10, 11, 12, 18, 19, 23, 26 and 27 are overexpressed in mammary tumors compared to normal mammary glands, but in addition these MaTARs are also present in comparable levels in at least one other tissue type. This information is crucial as we propose ASO-mediated knockdown of lncRNAs as a systemic treatment in breast cancer. ASOs cannot cross the blood-brain barrier; hence MaTARs that are also abundant in the brain (MaTAR1, 3, 7 and 27) should still be targetable in the tumor using systemic treatment without adverse effects on the brain.

To further characterize MaTARs in the context of global gene expression and to elucidate their potential as "driver genes" in mammary carcinogenesis, we performed weighted gene correlation network analysis (WGCNA) (Langfelder and Horvath 2008). We identified 39 modules with a median gene number of 274. Remarkably, 17 of the 30 MaTARs are residing within the same module. This particular module ("blue", Table S4) is enriched for genes that are highly expressed in our RNA-Seq datasets, both in the normal mammary gland organoids as well as mammary tumors (Figure S8B) and hence was termed the "mammary epithelium" module. It is rather large, comprising 5,744 genes in total. Interestingly, only 2,588 of the module genes are classified as "protein coding", emphasizing the tissue-specificity of non-coding RNA species as well as their potential functional role. In order to identify key drivers of the module, we focused on potential "hub genes" with high intra-module connectivity (> 1550) and very good correlation with the module eigengene (≥ 0.9), marked in red in Figure 3F. At least 6 MaTARs fall into this region, MaTAR5, 6, 9, 15, 16 and 30, indicating their importance within the module. We hypothesize that these MaTARs might be drivers of mammary carcinogenesis and/or mammary epithelium development. Notably, 16 of the 17 MaTARs in the "mammary epithelium" module were also highly expressed in tumor tissue compared to normal organs (Figure 3H). This indicates that MaTARs are excellent marker genes and sufficient to stratify mammary epithelia from other tissue types. A full list of all MaTARs and their network modules is provided in Table S4.

GO analysis of the module revealed association with several interesting biological processes. Most significant is the enrichment for the GO term "translation" along with "rRNA processing" and "ribosomal subunit biosynthesis". In addition, we detected "positive regulation of mRNA splicing via spliceosome", "regulation of cytokine production" and "immune response" as enriched GO terms. These findings are in excellent agreement with the top "hub genes" that include 33 ribosomal and snoRNA genes, several components of the spliceosome, cyclooxygenase 2 and 3 but also typical marker genes for mammary epithelia like caseins and lactalbumin. In terms of enriched cellular components, we detect "cytosolic ribosomal subunit", "extracellular exosome" and "focal adhesion" (Figure S8C). In summary, our findings demonstrate that we identified a module related to mammary epithelial development and/or mammary carcinogenesis, and suggest that MaTARs are likely to play key roles in these functional processes.

### Knockdown of MaTARs decreases cell viability and invasion

Based on the results of our computational approach, we proposed that MaTARs are driver genes in mammary cancer progression and/or metastasis and thus could potentially serve as novel therapeutic targets. We tested our hypothesis by performing knockdown experiments of all 30 MaTARs using antisense oligonucleotides (ASOs) in primary MMTV-PyMT tumor cells. These oligonucleotides are short (20-mers), single stranded DNA molecules containing phosphorothioate-modified nucleotides as well as modifications of the 2'-ribose (5-10-5 2'-MOE gapmer) (for review, see Geary et al. 2015). Upon binding of the ASO to the complementary target, the RNA-DNA duplex stimulates degradation of the lncRNA by RNase H and thereby reduces the steady-state level of the respective transcript (Wu et al. 2004). Importantly, we found that the uptake of ASOs in primary mammary tumor cells is efficient without the use of transfection agents, a mechanism that has been studied in detail in hepatocytes (Koller et al. 2011). While we tested all 30 MaTARs for ASO-mediated knockdown efficiency, one of the candidates, MaTAR2, was not expressed at levels high enough to be reliably quantified using qRT-PCR. This transcript was hence excluded from this experiment. For each of the MaTARs, we achieved knockdown efficiencies ranging from 38% - 89% after 24 h using 5 μM of the most potent ASO (Figure S9A).

To investigate the functional impact of MaTAR down-regulation on tumor cells, we combined the ASO treatment with cell viability or invasion assays (Figure 4A and B). Interestingly, MTT assays revealed that the knockdown of MaTAR12 leads to a 50% decrease in cell viability (Figure 4A), an effect comparable to ASO-mediated knockdown of Eg5 that served as a positive control (Koller et al. 2006). Four additional candidates, MaTAR6, 11, 18 and 20, show a less pronounced effect with a 20-30% decrease of cell viability. The remaining MaTARs did not seem to significantly impact tumor cell growth (< 20%). We did not observe an effect on cell viability with a scrambled ASO control (scASO) (Figure 4A). To confirm that our results obtained with the MTT assays are in fact detecting differences in cell viability, we performed fluorescence based cell viability assays using commercial kits on a subset of the MaTARs and obtained similar results (Figure S9B).

**Figure 4.**
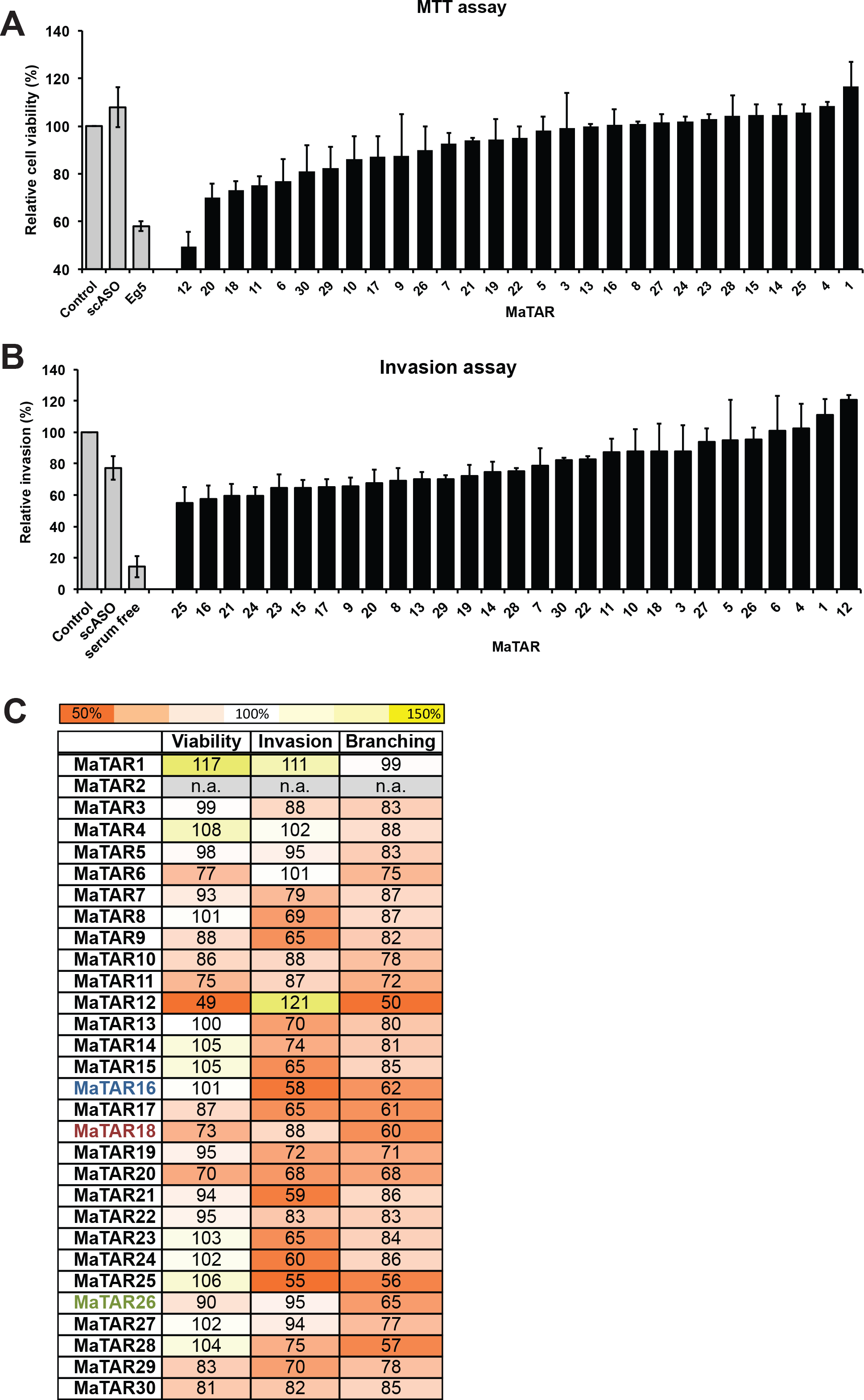
ASO-mediated knockdown of MaTARs in primary tumor cells. Primary mammary tumor cells were treated with 5 μM of the most potent ASO for 24 h. A) MTT assay after ASO-mediated knockdown of MaTARs. Cell viability is normalized to untreated cells ("Control"). A scrambled ASO ("scASO") was used as a negative control; ASO knockdown of Eg5 was used as a positive control. Bars denote the mean of two replicates +/- standard error. B) Invasion assay after ASO-mediated knockdown of MaTARs. Cell invasion is normalized to untreated cells ("Control"). A scrambled ASO ("scASO") was used as a negative control; serum-free medium was used as positive controls. Bars denote the mean of three replicates +/- standard error. C) Summary of observed effects on cell viability, cell invasion and organoid branching upon ASO-mediated knockdown of MaTARs. Data is normalized to untreated cells (= 100%). Reduction of viability, invasive potential or branching of > 20% compared to untreated cells was defined as a significant difference. The color legend is indicated.

In addition, we performed invasion assays of tumor cells in 96-well Boyden chamber plates upon ASO-mediated knockdown. We used serum-free medium as a positive control, as cells will not migrate through the basement membrane extract (BME) without a chemoattractant on the other side. We detected a 30-45% reduction of tumor cell invasion upon knockdown of MaTAR8, 9, 15, 16, 17, 20, 21, 23, 24 and 25 (Figure 4B). Furthermore, 20-30% reduction in invasion was observed for MaTAR7, 13, 14, 19, 28 and 29. While knockdown of almost all MaTARs impacts either the viability or the invasive potential of mammary tumor cells, indicating specific roles of the different non-coding transcripts in cellular processes, only MaTAR20 showed effects in both assays (Figure 4C). Of the 29 tested MaTARs nine (MaTAR1, 3, 4, 5, 10, 22, 26, 27 and 30) did not exhibit significant effects on cell growth in 2D or invasion upon knockdown. Our results indicate that our initial computational selection of up-regulated lncRNAs indeed identified a number of transcripts that have the potential to act as driver genes in tumor progression by impacting cell growth or invasion.

### Knockdown of MaTARs inhibits collective cell migration in organoids

To further elucidate the functional role of MaTARs in a more physiological context, we performed ASO-mediated knockdown experiments in mammary tumor organoids (Barcellos-Hoff et al. 1989; Fata et al. 2007; Ewald et al. 2008; Nguyen-Ngoc et al. 2012). As described above, organoids mimic aspects of the three-dimensional organization of an organ more closely than individual cells cultured in 2D. As for primary tumor cells, we found that mammary organoids perform transfection-free uptake of ASOs.

For the 29 MaTARs, we observed knockdown efficiencies in organoids ranging from 30 - 68% after 6 days of treatment using 4 μM of the most potent ASO (Figure S10A). Culturing the organoids for several days allowed us to observe any phenotypic changes that might be caused by the knockdown. Both cell proliferation and collective cell migration are required to form mammary organoid branches; therefore this experiment combines the readout of two functional assays. Untreated as well as scASO treated MMTV-PyMT organoids generally exhibit branching in about 70-75% of all organoids. Upon ASO-mediated knockdown, we detected a distinct decrease (30-50% less compared to untreated organoids) in duct formation for MaTAR12, 16, 17, 18, 20, 25, 26 and 28, as well as a mild decrease (20-30% less) for MaTAR6, 10, 11, 19, 27 and 29 (Figure S10B). The remaining 15 MaTARs did not show significant changes (< 20%) compared to untreated organoids. Interestingly, many of the MaTARs that interfered with cell viability and/or invasion in 2D assays also exhibited a branching defect in organoids (Figure 4C). For instance, the knockdown of MaTAR16, 17, 25 and 28 caused a very significant reduction of organoid branching of 38%, 39%, 44% and 43%, respectively (Figure S10A-D). All four transcripts also seemed to play a role in invasion in 2D culture, reducing invasion through an artificial BME by 42%, 35%, 45% and 25% (Figure 4B, C). In contrast, the down-regulation of MaTAR18 and MaTAR26 did not affect the invasive potential of tumor cells grown in 2D but reduced organoid branching significantly. MaTAR18 however did impair cell proliferation in 2D, which might contribute to the reduction of branching morphogenesis in 3D as well.

We further focused on MaTAR16 *(ENSMUSG00000086249, Gm12724),* MaTAR18 *(ENSMUSG00000085873, Ttc39aos1)* and MaTAR26 *(ENSMUSG00000097378, B230208H11Rik),* as these three transcripts showed very distinct organoid phenotypes upon ASO-mediated knockdown and had different affects on cell growth and invasion in 2D culture (Figure 4C). The three RNAs represent different transcript biotypes with MaTAR16 being classified as "processed transcript", MaTAR18 as "antisense" and MaTAR26 as "lincRNA". Both MaTAR16 and MaTAR26 are part of the "mammary epithelium" module, while MaTAR18 resides in another highly interesting module that is comprised of the gene-expression signature for embryonic stem cells and embryonic liver. This module is associated with GO terms related to replication and the cell cycle (data not shown).

Knockdown experiments were performed with two different ASOs for each transcript to ensure that the observed phenotypes are not caused by off-target effects. Phenotypes for the both ASOs are shown in comparison to organoids that were either untreated or treated with scASO (Figure 5A with additional images in Figure S10C). While knockdown of MaTAR16 and MaTAR26 completely abolished organoid branching, down-regulation of MaTAR18 resulted in organoids without defined branches but with a "rough" surface. In addition, we observed that organoids treated with ASOs targeting MaTAR16 tend to be smaller compared to untreated or scASO treated organoids (Figure 5A). Individual knockdown efficiencies are displayed in Figure 5B and correlate well with the quantitation of branching (Figure 5C). Compared to untreated controls, we detected a reduction in branching of 38% for MaTAR16, 40% for MaTAR18 and 35% for MaTAR26 with the best ASO, respectively.

**Figure 5.**
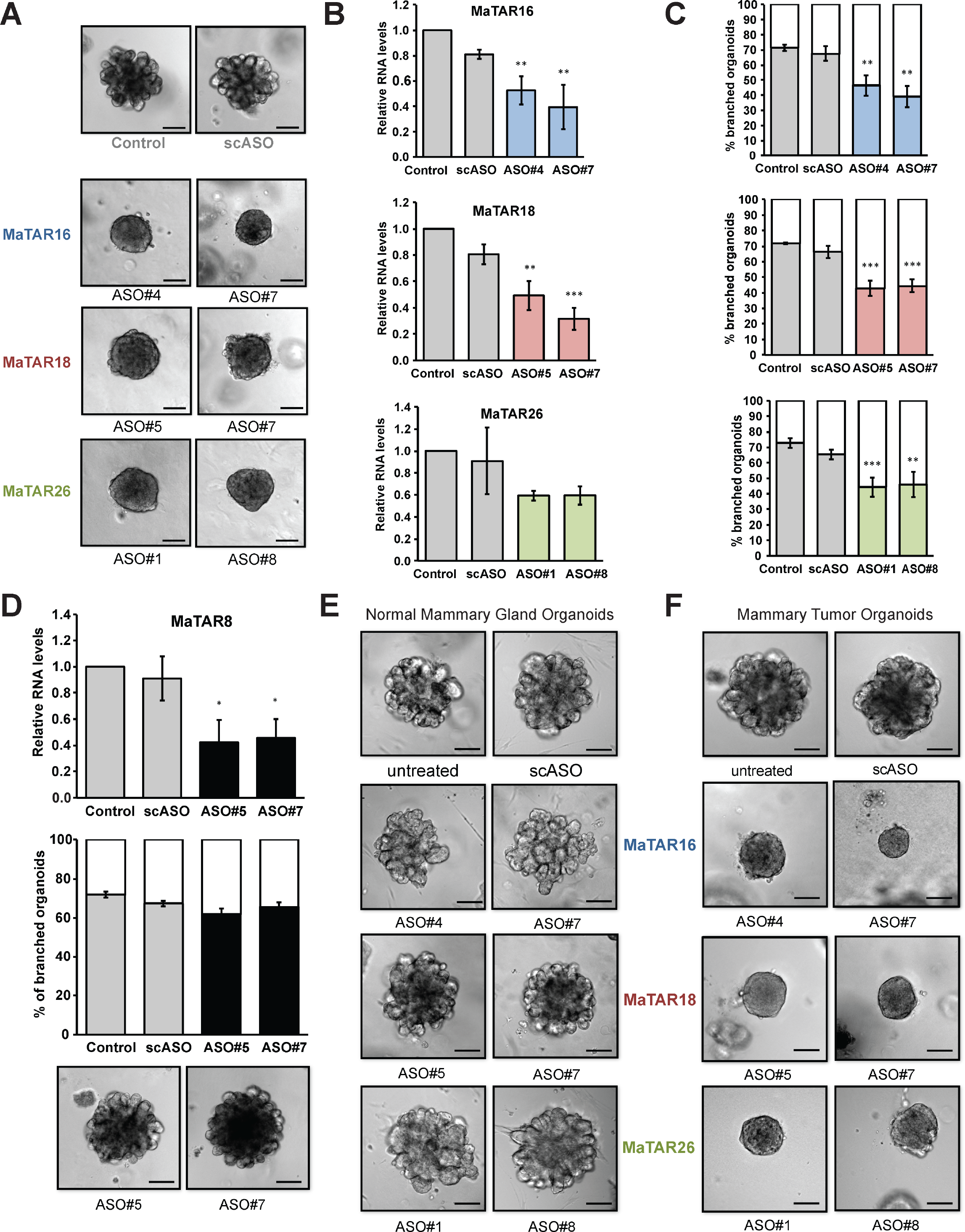
ASO-mediated knockdown of MaTARs in mammary organoids. Organoids were treated with 4 μM of specific ASOs for 6 days. Statistical significance was determined using a two-tailed, paired student's t-test; *p < 0.05, ** p < 0.01, *** p < 0.01. A) DIC images of tumor organoids upon knockdown with the two most potent ASOs targeting MaTAR16, MaTAR18 or MaTAR26 are shown. Knockdown of MaTAR16 and MaTAR26 leads to a loss of branching while knockdown of MaTAR18 results in organoids with tiny protrusions. scASO = scrambled ASO control. Scale bar = 100 μm. B) qRT-PCR to determine the knockdown efficiency of three specific ASOs per MaTAR, normalized to untreated cells ("Control"). Bars denote the mean of three replicates +/- standard deviation. C) Quantification of tumor organoid branching upon knockdown with three specific ASOs per MaTAR. Reduction of organoid branching correlates directly with ASO-mediated knockdown efficiency. Black and grey bars indicate the percentage of branched organoids +/- standard deviation; white bars indicate the percentage of unbranched organoids. The mean of three biological replicates is shown; total number of assayed organoids per treatment = 300. D) Knockdown of MaTAR8 using two specific ASOs in tumor organoids. Despite efficient MaTAR8 knockdown, no phenotypic changes or reduction of organoid branching was observed. Upper panel: qRT-PCR to determine the knockdown efficiency. Bars denote the mean of three replicates +/- standard deviation. Middle panel: Quantification of organoid branching. The mean of three biological replicates is shown; total number of assayed organoids per treatment = 300. Lower panel: DIC images of organoids upon knockdown. Scale bar = 100 μm. Control = untreated organoids, scASO = scrambled ASO control. E) DIC images of WT normal mammary gland organoids upon knockdown with the two most potent ASOs targeting MaTAR16, MaTAR18 or MaTAR26 are shown. No change in organoid branching was observed upon MaTAR knockdown. scASO = scrambled ASO control. Scale bar = 100 μm. F) DIC images of C57Bl/6 MMTV-PyMT organoids upon knockdown with the two most potent ASOs targeting MaTAR16, MaTAR18 or MaTAR26 are shown. Phenotypic changes resemble the reduction of organoid branching in A). scASO = scrambled ASO control. Scale bar = 100 μm. **Figure 6**

As expected, we do not see phenotypic changes upon ASO-mediated down-regulation for all MaTARs. While we obtained a knockdown efficiency of 55% for MaTAR8 *(ENSMUSG00000092569, Gm20544, lncRNA-Smad7),* we did not observe a loss of branching or any other phenotypic change in organoids (Figure 5D, additional images in Figure S10D). This is in agreement with previous findings describing that MaTAR8 is not involved in epithelial mesenchymal transition (EMT) (Arase et al. 2014). Furthermore, MaTAR7 and MaTAR13 did not change branching morphogenesis despite a knockdown efficiency of > 50% (Figure S10A, B, D) and promising invasion assay results for MaTAR13 (Figure 4B-C). These results indicate that different MaTARs impact different aspects of tumor biology.

To further support the role of MaTAR16, MaTAR18 and MaTAR26 as excellent therapeutic targets, we performed ASO-mediated knockdown in control organoids derived from WT nulliparous mammary glands. Knockdown efficiencies of the ASOs are comparable to the experiments in tumor organoids (Figure S11A). Importantly, we do not see a pronounced loss of branching in the normal mammary gland organoids, strongly indicating the cancer-specific role of these three lncRNAs (Figure 5E).

To exclude the possibility that our results are dependent on the genomic background of the mice, we confirmed the findings for MaTAR16, MaTAR18 and MaTAR26 in organoids derived from C57Bl/6 MMTV-PyMT mice. In fact, we observe very similar phenotypes for MMTV-PyMT organoids generated from both FVB and C57Bl/6 backgrounds (Figure 5F). Knockdown efficiencies and branching quantifications are shown in Figure S11B.

### Human MaTARs are amplified and overexpressed in breast cancer

We identified and characterized MaTARs in mouse models of human breast cancer and propose that several MaTARs act as key drivers of mammary tumor cell growth and/or migration. To confirm the relevance of these lncRNAs in human breast cancer we set out to identify human counterparts of all 30 MaTARs. Therefore, we compared mouse and human transcripts on the level of both sequence conservation and genomic location. First, we extracted the mouse MaTAR sequence and screened for transcripts with sequence overlap in the human genome using the BLAT alignment tool. If the mouse MaTAR sequence overlapped significantly with an annotated human transcript, this transcript was defined as the human counterpart. For instance, BLAT results for MaTAR1 indicate sequence identify in the human genome in a similar context, mapping to the last exon of a longer lncRNA, LINC00461, and in close spatial proximity of a miRNA gene (Figure 6A). The sequence of MaTAR1 spans 2419 bp in the human genome with a sequence identity of 92.6% and a BLAT score of 1438. This genomic region is highly conserved in vertebrates as indicated by the PhyloP conservation track (Figure 6A). For some MaTARs, similarly high sequence identities were obtained while we did not observe any sequence overlap for other candidates. This sequence-based approach is not informative for pseudogenes, as the parental protein-coding genes are the best sequence matches, e.g. *FAM96B* for a *Fam96b* pseudogene such as MaTAR2.

**Figure 6.**
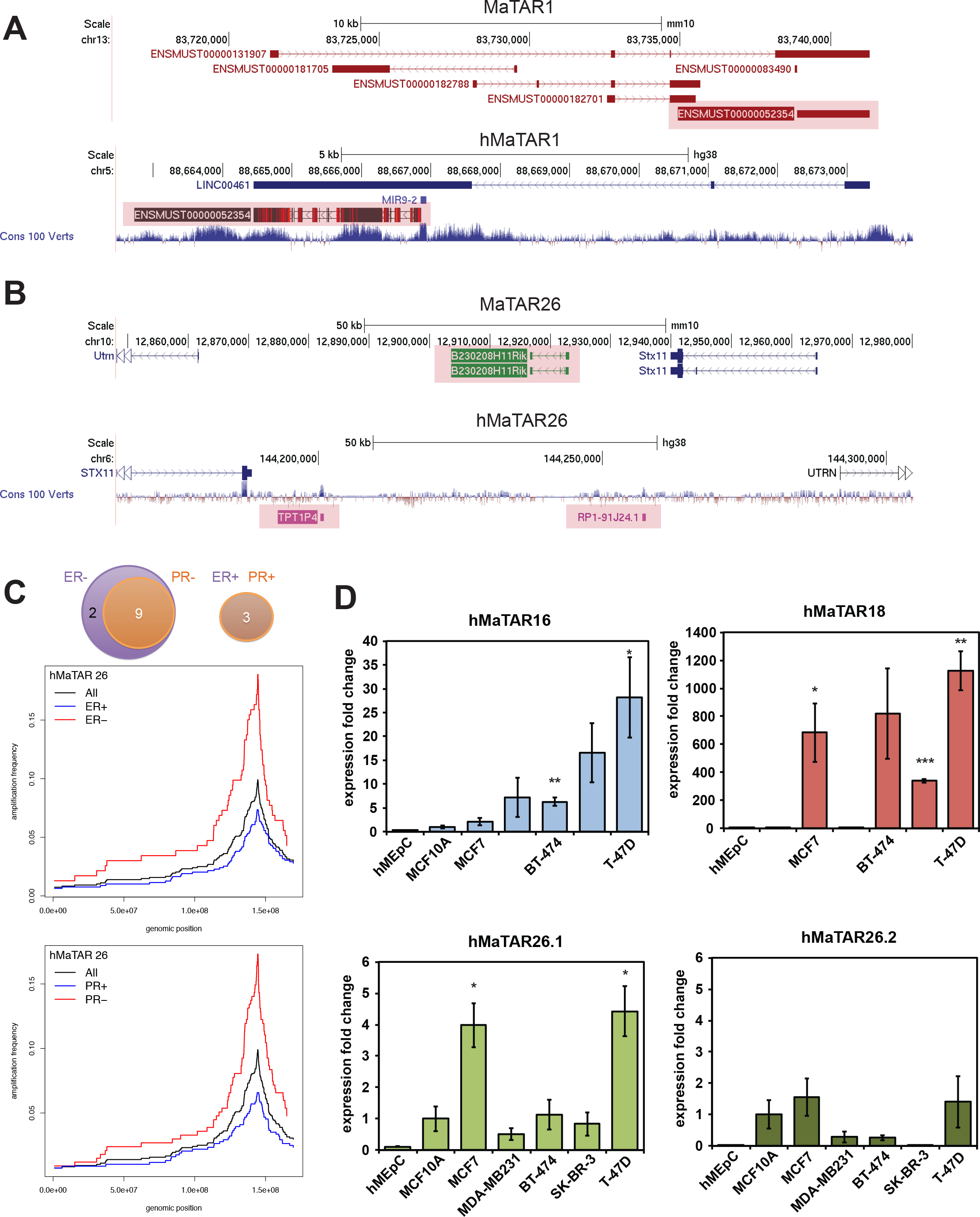
hMaTARs are amplified and overexpressed in breast cancer. A) Identification of a human MaTAR gene based on sequence identity. The upper panel represents the mouse genome (mm10), the lower panel the human genome (hg38). MaTAR1 is highlighted (ENSMUST00000052354). B) Identification of a human MaTAR gene based on genomic location (synteny). The upper panel represents the mouse genome (mm10), the lower panel the human genome (hg38). C) Upper panel: Venn diagram illustrating hMaTARs significantly amplified in ER-negative and/or PR-negative tumors (left) as well as in ER-and PR-positive tumors (right). Middle and lower panel: Amplification frequency profiles for hMaTAR26 (location) in TCGA breast cancer (BRCA) data set for all 1080 samples in the set and separately within the subsets with a given hormone receptor status. D) qRT-PCR of hMaTAR16, 18 and 26 in human breast cancer cell lines and normal mammary epithelial cells. Expression is normalized to MCF10A cells. Bars indicate the mean of three independent biological replicates for all cells but hMEpC (n=2), error bars indicate the standard error. A two-tailed, paired student's t-test was performed; * p < 0.05, ** p < 0.01, *** p< 0.001.

As many non-coding RNAs are conserved between different species on the level of genomic location rather than based on sequence, possibly implying functional conservation (Diederichs 2014; Ulitsky et al. 2011), we extended our analysis to synteny by analyzing the nearest neighboring genes of each MaTAR. If the genomic location of a MaTAR is in close proximity of a protein-coding gene, we screened the surroundings of the same protein-coding gene in the human genome. For instance, the MaTAR26 gene is located between the two protein coding genes, *Utrn* and *Stx11* (Fig 6B). There are two annotated non-coding genes between *STX11* and *UTRN* in the human genome as well, *TPT1P4* and *RP1-91J24.1*. We identified human counterparts for all MaTARs based on sequence or synteny. In several cases, we observed conservation on both sequence and synteny level. The complete list of human MaTARs (hMaTARs) including genomic locations is provided in Table S5. Interestingly, many MaTARs are located in clusters of two or more non-coding genes in the genome. Hence, to identify the correct human counterpart unambiguously, experimental validation is necessary.

To validate that the identified hMaTARs are clinically relevant, we performed extensive analysis of breast cancer data derived from The Cancer Genome Atlas (TCGA). Interestingly, loci containing 14 hMaTARs are amplified significantly in more than 10% in breast cancer across all breast tumors (Table S6). While we did not detect correlations of hMaTAR locus amplification and the expression subtypes determined by PAM50, we identified significant associations between hMaTAR amplification and the status of the hormone receptors as measured by IHC. Interestingly, most hMaTARs are more often amplified in the ER-negative and/or PR-negative patients than hormone receptor positive patients (Figure 6C, Table S6). 9 hMaTARs are amplified in ER-and PR-negative tumors and 2 hMaTARs amplified in only ER-negative breast tumors. Three additional hMaTARs are amplified specifically in ER-and PR-positive tumors (Figure 6C, Table S6).

Both hMaTAR16 and hMaTAR18 are amplified in about 7.5% of all breast tumors independently of the ER/PR status (p-value > 0.05, Table S6). Notably, hMaTAR26 is amplified in about 10% of all tumors and correlates significantly (p < 0.05) with both ER and PR status. In fact, the hMaTAR26 locus is amplified in almost 20% of all ER and PR negative tumors (Figure 6C, Table S6).

To investigate whether hMaTARs exhibit copy number gains only or are in fact overexpressed in breast cancer compared to normal mammary gland cells, we performed qRT-PCR of hMaTAR16, 18 and 28 in five different human breast cancer cell lines as well as MCF10A and primary hMEpC cells, representing the untransformed mammary epithelium. Notably, all three hMaTARs are overexpressed significantly in one or more human breast cancer cell lines (Figure 6D). Individual hMaTARs are preferentially overexpressed in some cell lines versus others. In the case of MaTAR26, two non-coding RNAs were identified as human counterparts based on synteny (Figure 6B, Table S5). The expression analysis clearly indicates that hMaTAR26.1 *(TPT1P4)* rather then hMaTAR26.2 *(RP1-91J24.1)* is overexpressed in breast cancer (Figure 6D). As hMaTAr26.2 and hMaTAR26.1 are in close spatial proximity, our results indicate that while the whole genomic locus is amplified (Table S6) *TPT1P4* is up-regulated specifically.

In summary, we performed an RNA-Seq screen to identify lncRNAs that are up-regulated in mammary tumors and represent potentially oncogenic transcripts. We identified 109 lncRNAs that are up-regulated at least 2-fold in MMTV-PyMT and 207 that are up-regulated at least 2-fold in MMTV-Neu-NDL tumors. Among these transcripts we prioritized 30 as MaTARs and we identified human orthologs. We showed that 15 of the 30 MaTARs that were functionally validated are involved in cell proliferation and/or migration processes. ASO-mediated knockdown of these transcripts leads to a reduction or complete loss of organoid branching. This effect is specific only for mammary tumor-derived organoids but not normal mammary gland-derived organoids, indicating that MaTARs represent promising therapeutic targets in breast cancer.

## Discussion

In recent years, high-throughput sequencing studies have begun to shed light on the emerging role of non-coding transcripts in cancer (for review, see Prensner and Chinnaiyan 2011; Wapinski and Chang 2011; Shore et al. 2012; Wahlestedt 2013; Cheetham et al. 2013). However, many global gene expression studies were performed in human cancer cell lines. Studying tumor biology in cell lines is a practical but overly simple approach. There are several potential caveats to using cell lines such as adaption to growth *in vitro* on plastic dishes, lack of corresponding normal tissue controls, cell immortalization, lack of a physiological ECM and accumulation of chromosomal aberrations due to continued passaging. Importantly, transcriptome analyses revealed that commonly used cancer cell lines often do not represent tumors well enough and that tumors cluster closer to their corresponding normal tissues than to cell lines (van Staveren et al. 2009; Domcke et al. 2013). Primary 3D *ex vivo* organoids resemble the organization and physiology of native epithelia much better than cancer cell lines grown in 2D, and additionally model interactions with the ECM (Boj et al. 2015; for review, see Sachs and Clevers 2014; Shamir and Ewald 2014). To investigate whether organoids mirror the tumor on the level of global gene expression as well, we performed the first RNA-Seq screen comparing the transcriptome of organoids to the mammary tumors from which they were derived. Importantly, our data demonstrate that the vast majority of genes (99% of the transcriptome) do not significantly change in expression when comparing three replicates of primary tumor sections to organoids cultured for 6 days in matrigel. Importantly, non-coding RNA levels seem to remain virtually unchanged. Our findings support the potential use of patient-derived organoids as a model system to study tumor biology and to model drug responses ("personalized medicine") (Boj et al. 2015; for review, see Shamir and Ewald 2014; Lancaster and Knoblich 2014).

Genome-wide gene expression studies are helpful resources but often lack systematic functional validation. Here, we further characterized the newly identified MaTARs using both computational approaches and molecular assays. Motif analysis of all differentially regulated lncRNAs in our study revealed specific regulators for up-and down-regulated non-coding transcripts. Repressed lncRNAs show an enrichment for the transcription factor Kaiso, which has been described to act as an oncogene by DNA methylation-dependent silencing of tumor suppressor genes in colon cancer (Lopes et al. 2008). Furthermore, we detect an enrichment of motifs for NF-_K_B and CEBP, two transcription factors that have been implicated in aggressive breast cancer and epithelial mesenchymal transition (EMT) (Huber et al. 2004; Bundy and Sealy 2003). Up-regulated lncRNA promoters are enriched for AR and p63. AR was described to stimulate breast tumor progression but its effect depends on the ER status of the patient (for review, see Chang et al. 2013). The transcription factor p63 is required to initiate collective cell migration and invasion in tumor organoids and *in vivo* (Cheung et al. 2013). In summary, our motif analysis revealed that the same regulators that control known tumor suppressor genes might down-regulate the identified lncRNAs, while up-regulated lncRNAs are activated by factors that also impact oncogenes. In both cases, the key regulators are associated with EMT and metastasis, indicating that the differential expression of non-coding RNAs correlates with tumor progression.

We applied weighted gene correlation network analysis (Langfelder and Horvath 2008) to explore gene sets that are co-expressed with MaTARs. Seventeen of the 30 MaTARs fell into the "mammary epithelium" module, with MaTAR5, 6, 9, 15, 16 and 30 standing out due to high intra-module connectivity and eigengene correlation. These MaTARs are likely key drivers of mammary epithelium development and/or mammary carcinogenesis. Interestingly, the GO terms associated with this network module show a strong enrichment for ribosome biogenesis. A cluster of co-expressed ribosomal protein genes has been observed previously in normal human breast samples and breast tumors (Perou et al. 1999). Furthermore, several studies described alterations of ribosome biogenesis components in breast cancer cells, such as elevated snoRNAs overexpression and alterations in rRNA modifications (Belin et al. 2009; Su et al. 2013). In addition, the nuclear Erbb2/neu protein was described to interact with RNA Polymerase I and thereby stimulate rRNA transcription and cell growth in breast cancer cells (Li et al. 2011). Hence, we conclude that high expression of ribosome biogenesis factors likely demarcates the normal and/or malignant mammary epithelium specifically. Additional GO terms associated with the "mammary epithelium" module are the spliceosome, exosomes and focal adhesions. Core components of the spliceosome were described to be overexpressed in several cancers and represent the highest enriched cluster when comparing aggressive breast cancer to benign lesions (André et al. 2009; Bonnal et al. 2012; Anczuków et al. 2015). Both exosomes and focal adhesions, one of the main types of adhesion in epithelia, play an important role in mammary gland development and breast cancer (for review, see Hendrix and Hume 2011; Schedin and Keely 2011). Taken together, the GO terms enriched in this module functionally characterize it as specific for the mammary epithelium. In addition to many MaTARs, this module contains hundreds of non-coding RNAs such as lincRNAs, pseudogenes or miRNA host genes, indicating that these RNA species likely play an important role in malignant and/or normal mammary epithelial growth processes.

Upon further functional validation using ASO-mediated knockdown experiments in primary mammary tumor cells and mammary organoids we observed that down-regulation of many MaTARs leads to a reduction of cell proliferation, invasion and/or collective cell migration, observed as loss of organoid branching. Specifically, knockdown of MaTAR16, MaTAR18 and MaTAR26 significantly decreased organoid branching in a cancer-specific context. Taken together, our functional validation confirmed that MaTARs are potential drivers of tumor cell proliferation and/or collective cell migration.

Based upon our genome-wide analysis of mouse models that recapitulate two different sub-types of human breast cancer we propose that the human counterparts of MaTARs are likely important therapeutic targets. In this regard, concerns are often raised about the lack of conservation between mouse and human lncRNAs (Derrien et al. 2012). Unlike protein-coding genes, the functions of lncRNAs are not necessarily directly linked to their nucleotide sequence but are often dependent on RNA secondary structure folding or syntenic transcription (Diederichs 2014). Synteny between non-coding and protein-coding genes is frequently conserved across species (Ulitsky et al. 2011). Importantly, we identified human counterparts for all MaTARs and showed that many of these genes are amplified in ER-negative and/or PR-negative breast cancer patients (TCGA data) and overexpressed in breast cancer cell lines.

Down-regulation of MaTARs using an ASO-based treatment might be a viable option of reverting the transcript levels back to those of a normal mammary epithelium, thereby negatively impacting tumor progression. We propose ASO-mediated knockdown of MaTARs as an exciting new treatment strategy in breast cancer. Ongoing clinical trials recently demonstrated that systemic administration of ASOs is a safe and effective therapy (Büller et al. 2015). Like most lncRNAs, MaTARs are expressed tissue-or tumor-specifically, hence systemic treatment with ASOs is unlikely to exhibit significant effects on other normal tissues where the drug target is virtually absent.

Understanding how MaTARs influence proliferation and collective cell migration in a cancer-specific context is essential to validate them as key drivers of mammary tumor initiation, progression and/or metastasis. Future studies will focus on elucidating the molecular mechanism by which MaTARs impact cellular function in cancer.

## Methods

### Animals

Animal experiments were carried out in the CSHL Laboratory Animal Shared Resource, in accordance with IACUC approved procedures. MMTV-PyMT mice (Guy et al. 1992) were obtained from Dr. Mikala Egeblad (CSHL). MMTV-Neu-NDL mice (Siegel et al. 1999) were obtained from Dr. Senthil Muthuswamy (CSHL). MMTV-Cre and Flox-Neo-Neu-NT mice as well as tumors derived from MMTV-Cre;Flox-Neo-Neu-NT mice (Andrechek et al. 2000) were obtained from Dr. William Muller (McGill University, Canada). All mouse strains are in the FVB/N background. Wild type FVB/N mice were ordered from The Jackson Laboratory for control experiments. Tumors and normal mammary glands were extracted immediately after euthanizing the animal and processed to generate primary cells, organoids or tissue sections.

### Organoid culture

All cell culture reagents were obtained from Gibco (Life Technologies) unless stated otherwise. Organoids from wild type nulliparous mammary glands, MMTV-PyMT tumors and MMTV-Neu-NDL tumors were prepared and cultured as described previously (Ewald 2013). For MMTV-PyMT mice, individual tumors were isolated and organoids were generated in a tumor-specific manner. Mammary epithelial fragments (organoids) were mixed with growth factor-reduced matrigel (Corning) at a concentration of 5 organoids/μl and plated as 80 μl domes in Cellstar 24-well dishes (Greiner Bio One). Organoids were grown in DMEM/F12 medium supplemented with 1x ITS (insulin, transferrin, and sodium selenite) media supplement (Sigma), 1% penicillin/streptomycin and 2.5 nM murine FGF2 (Peprotech).

### Primary cell culture

All cell culture reagents were obtained from Gibco (Life Technologies) unless stated otherwise. To generate primary mammary tumor cells, the protocol for organoid preparation was followed by additional steps to dissociate organoids into single cell suspensions. Organoids were treated with 0.25% Trypsin and 0.1% EDTA in PBS for 5 min at 37°C. To stop the reaction, 10 ml DMEM/F12 medium supplemented with 10% FCS and 4 U/μl of DNase I (Sigma) was added and incubated for 15 min at 100 rpm and 37°C. The cell solution was spun down for 10 min at 520 g. The pellet was resuspended in 10 ml DMEM/F12 with 10% FCS and 1% penicillin/streptomycin and filtered via a 70 μm cell strainer. Three additional washing steps (centrifuge for 5 min at 520 g, resuspend the pellet in 10 ml DMEM/F12 with 10% FCS and 1% penicillin/streptomycin) were performed. The final cell pellet was resuspended and cultured in DMEM/F12 medium supplemented with 10% FCS, 1% penicillin/streptomycin and 5 mg/ml insulin (Sigma).

### RNA-Sequencing

Total RNA was isolated either directly from cryosections of the tumor tissue or from organotypic epithelial cultures using TRIzol (Life Technologies) according to the manufacturer's instructions. For tissue sections, the tumors were embedded in OCT and cryosectioned. Sections from the middle of the tumor were stained using toluidine blue (Sigma) and assayed regarding the homogeneity of the section. Homogenous 30 μm sections comprising >90% malignant cells were immediately dispersed and homogenized in TRIzol. RNA quality was assayed by running an RNA 6000 Nano chip on a 2100 Bioanalyzer (Agilent). For high-throughput sequencing, RNA samples were required to have an RNA integrity number (RIN) ≥ 9. TruSeq (Illumina) libraries for polyA+ RNA-Seq were prepared from 0.5 - 1 μg RNA per sample. To ensure efficient cluster generation, an additional gel purification step of the libraries was applied. The libraries were multiplexed (4-6 libraries per lane) and sequenced paired-end 101 bp on the HiSeq2000 platform (Illumina), resulting in on average 40 Mio reads per library.

### Computational analysis

The quality of the raw data was evaluated using FastQC (http://www.bioinformatics.babraham.ac.uk/projects/fastqc/), and reads were mapped to mm10 using STAR v2.4.1 (Dobin et al. 2012), resulting in an overall mapping efficiency of >90%. The GENCODE mV5 GTF was used as a reference and the reads per gene record were counted using the HTSeq package v0.5.4p5 (Anders et al. 2015) and parameters-m union-s no. Differential gene expression was performed with DESeq2 v1.8.1 (Love et al. 2014). An adjusted p-value of < 0.1 was set as threshold for statistical significance of differential gene expression. Functional analysis of KEGG pathways was carried out using the R/Bioconductor packages GAGE (Luo et al. 2009) and Pathview (Luo and Brouwer 2013). Gene set enrichment analysis was performed using the javaGSEA desktop application (Subramanian et al. 2005). Gene ontology (GO) analysis was carried out using GOrilla (Eden et al. 2009) separately on genes up-or down-regulated compared to normal mammary gland organoids. The coding potential was calculated with CPAT (Wang et al. 2013). Motif enrichment at promoter regions was performed using HOMER (Heinz et al. 2010).

### Weighted gene coexpression network analysis

Weighted gene coexpression network analysis (WGCNA) was carried out as described previously (Bergmann et al. 2015). Briefly, ENCODE data sets (CSHL long RNA sequencing), ESC data (Bergmann et al. 2015) as well as the tumor and organoid RNA-Seq datasets were used as input. We performed variance-stabilizing transformation of HTSeq-generated counts and averaged the counts from replicate samples. The WGCNA R package (Langfelder and Horvath 2008) was used with a value of β = 5 as empirically chosen soft-threshold. This analysis identified 39 modules with a median gene number of 274.

### ASO-mediated knockdown in primary cells

MMTV-PyMT primary cells were seeded at a density of 20,000 cells/well (100 μl per well) into 96-well plates. Transfection-free uptake of ASOs was accomplished by adding 5 μM of either a MaTAR-specific ASO or scrambled ASO (scASO) to the primary cell culture medium immediately after seeding the cells. ASO sequences are provided in Supplementary Table S5. Cells were incubated for 24 h at 37°C and RNA was isolated using the RNeasy 96 kit (Qiagen) according to the manufacturer's instructions. RNA samples were used directly in a one-step 384-well qRT-PCR (QuantiTect SYBR Green RT-PCR Kit, Quiagen) on an Applied Biosystems 7900HT Fast Real-Time PCR System (Thermo Fisher Scientific). qRT-PCR conditions were as follows: 30 min at 50°C for reverse transcription, 15 min at 95°C for the initial activation step followed by 40 cycles of 15 sec at 94°C, 30 sec at 60°C. *Peptidylprolyl isomerase B (cyclophilin B)* was used as an endogenous control to normalize each sample, and relative expression results were calculated using the 2^-ΔΔCt^ method. A list of primers used is provided in Supplementary Table S6.

### ASO-mediated knockdown in organoids

Organoids were seeded at a density of 5 organoids/μl and plated as 80 μl domes in Cellstar 24-well dishes (Greiner Bio One). Transfection-free uptake of ASOs was accomplished by adding 4 μM of either a MaTAR-specific ASO or scASO to the organoid culture medium 15-20 min after the organoids were plated in matrigel domes. Organoids were incubated for 6 days at 37°C and both medium and ASOs were replenished at day 3. ASO sequences are provided in Supplementary Table S5. Total RNA was isolated using TRIzol (Life Technologies). DNase I (Life Technologies) treatment was performed for 15 min at RT to remove contaminating DNA. cDNA synthesis was carried out using TaqMan Reverse Transcription reagents and random hexamers (Life Technologies) according to the manufacturer's instructions. Real time quantitative PCR was performed using the Power SYBR Green Master Mix (Life Technologies) in 384-well plates using the Applied Biosystems 7900HT Fast Real-Time PCR System (Thermo Fisher Scientific). Cycling conditions were as follows: 10 min at 95°C followed by 40 cycles of 15 sec at 95°C, 1 min at 60°C. Primer specificity was initially tested by agarose gel electrophoresis and subsequently monitored by melting curve analysis. *Peptidylprolyl isomerase B (cyclophilin B)* was used as an endogenous control to normalize each sample and relative expression results were calculated using the 2^-ΔΔCt^ method. A list of primers used is provided in Supplementary Table S6. For visualization purposes and quantification of organoid branching, images were acquired using an Observer Live Cell inverted microscope (Zeiss) at 10x magnification.

### Cell viability assays

MMTV-PyMT primary tumor cells were seeded at a density of 10,000 cells/well (100 per well) into 96-well plates and treated with 5 μM of either a MaTAR-specific ASO or scASO. Cells were grown for 72 h at 37°C. 10 μl MTT solution (Cell Growth Determination Kit, MTT based; Sigma) was added to the wells and incubated for 4 h at 37°C. Next, 100 μl MTT solvent was added directly to the wells to ensure total solubility of the formazan crystals and incubated for 10 min with shaking. Measurements of absorbance at 570 μm were performed using a SpectraMax i3 Multi-Mode Detection Platform (Molecular Devices). Background absorbance at 690 nm was subtracted. Alternatively, CellTiter-Fluor™ Reagent (Promega) was added, incubated for 30 min at 37 °C and fluorescence was detected using the 400/505 nm filter set. Percent viability in the ASO treated cells was calculated as a percentage of the mock treated control.

### Invasion assay

The invasive potential of MMTV-PyMT primary tumor cells was assessed using Cultrex^®^ 96 well BME Cell Invasion Assay (Trevigen). Cells were starved in DMEM/F12 medium containing 0.1% FBS for 24 h, then harvested and seeded at a density of 5 x 104 cells/well into the invasion chamber. 5 μM MaTAR-specific ASO or scASO were added to 150 μl of growth medium containing 10% FBS. As a negative control, serum-free medium was used that did not stimulate cell invasion through the BME. The plate was incubated at 37°C for 24 h and the assay was performed according to the manufacturer's instructions. Briefly, the tumor cells that invaded through the BME layer and attached to the bottom of the invasion chamber were collected using cell dissociation solution and stained with Calcein AM solution. The fluorescence was measured with a SpectraMax i3 Multi-Mode Detection Platform (Molecular Devices) using the 480/520 nm filter set. Each sample was measured in triplicate.

### Expression analysis of human breast cancer cell lines

RNA from Human Mammary Epithelial Cells was obtained from Cell Applications, Inc. MCF10a cells were cultured in DMEM/F12 media, supplemented with 5% horse serum (Invitrogen), 0.5 μg/ml hydrocortisone, 10 μg/ml insulin, 0.1 μg/ml cholera toxin and 20 ng/ml EGF. MCF7 cells were cultured in DMEM media (Corning) supplemented with 20% FBS and 0.1 mM MEM Non-Essential Amino Acids (Gibco). MDA-MB 231 cells were cultured in DMEM media (Corning) supplemented with 10% FBS. BT-474 cells were cultured in Improved MEM (Gibco) supplemented with 10% FBS and 10 ug/ml insulin (Sigma). SK-BR-3 cells were cultured in McCoy's 5a Media (Corning) supplemented with 10% FBS. T-47D cells were cultured in RPMI-1640 Media (Corning) supplemented with 10% FBS. Cells were placed in a 37°C incubator at 5% CO2, and total RNA was isolated using TRIzol reagent (Ambion, Life Technologies) according to the manufacturer's instructions. Synthesis of cDNA and qRT-PCR was performed as described for organoids (above).

### Somatic DNA copy number alterations in human breast cancer

DNA copy number status of all hMaTAR candidate loci was examined in the set of 1080 segmented copy number profiles of breast tumors collected by TCGA (http://cancergenome.nih.gov/). Specifically, for a given profile and locus *r*_min_, the lowest *r ≡* log_2_(tumor to matching normal copy number ratio) anywhere within the locus was determined, along with a segment *s* overlapping the locus with this value of *r.* The covering copy number event for the locus was defined as the largest contiguous interval containing s such that *r* ≥ *r*_min_ everywhere within the interval. Loci with *r*_min_ ≥ 0.2 were deemed amplified. Based on this definition, a number of descriptive statistical quantities were computed for each locus, as specified in Table S6. These are the overall amplification frequency; the frequency of covering events shorter than 5Mb; the frequency of covering events shorter than 10Mb; the amplification frequencies separately for each value (positive or negative) of the hormone (estrogen or progesterone) receptor status (ER±, PR±). Amplification frequency profile for each locus was determined by computing, for each genomic position in the chromosome containing the locus, the frequency of amplified covering events at that position. Statistical tests were performed for association of amplification at each locus with the status of each of the hormone receptors, with the results listed in Table S6. Specifically, Fisher exact tests were conducted for association of the hormone receptor status with all amplifications at the locus, and the corresponding p-values and odds ratios were tabulated. The analysis was then repeated including only amplifications with covering events shorter than 10Mb, and the results likewise tabulated. Finally, additional analysis of ER (PR) association was performed to adjust for genome-wide genomic instability. To this end, the amplified fraction *F_P_* of the genome was computed for each copy-number profile *P,* and the distribution of the quantity *A_LP_ = I_LP_* - *F_P_* was examined, where *I_LP_* is the (0,1)-valued indicator for amplification at locus *L* in profile *P. A_LP_*is therefore a mean-subtracted (centered) indicator for amplification at locus *L* in profile *P.* Non-parametric Wilcoxon rank test for association with the ER (PR) was conducted for this variable at each locus and the corresponding p-values listed in Table S6.

### Data Access

The RNA-Seq data presented in this paper are publicly available at the NCBI Gene Expression Omnibus (GEO; http://www.ncbi.nlm.nih.gov/geo) under accession number GSE72823.

## Acknowledgements

We would like to thank all members of the Spector Lab for helpful discussions and suggestions throughout the course of this study, in particular Gayatri Arun for critically reviewing the manuscript. We acknowledge Mikala Egeblad, Senthil Muthuswamy (University of Toronto) and William J. Muller (McGill University) for providing mice and/or tumors. We acknowledge the CSHL Animal Shared Rresource, Animal and Tissue Imaging Shared Resource, DNA Sequencing Shared Resource, and Microscopy Shared Resource (NCI 2P3OCA45508). The Manhasset Women's Coalition Against Breast Cancer (S.D.D.) and the NCI 5P01CA013106-Project 3 (D.L.S.) supported this research.

## Author Contributions

S.D.D. and D.L.S. conceived the study, designed the experiments, and wrote the manuscript. S.D.D. performed the computational analyses. S.D.D. and K-C.C. performed experiments and interpreted the data. S.M.F., F.R. and C.F.B. designed and provided ASOs. J.S. and A.K. analyzed TCGA data.

